# A Deep SE(3)-Equivariant Model for Learning Inverse Protein Folding

**DOI:** 10.1101/2022.04.15.488492

**Authors:** Matt McPartlon, Ben Lai, Jinbo Xu

## Abstract

In this work, we establish a framework to tackle the inverse protein design problem; the task of predicting a protein’s primary sequence given its backbone conformation. To this end, we develop a generative SE(3)-equivariant model which significantly improves upon existing autoregressive methods. Conditioned on backbone structure, and trained with our novel partial masking scheme and side-chain conformation loss, we achieve state-of-the-art native sequence recovery on structurally independent CASP13, CASP14, CATH4.2, and TS50 test sets. On top of accurately recovering native sequences, we demonstrate that our model captures functional aspects of the underlying protein by accurately predicting the effects of point mutations through testing on Deep Mutational Scanning datasets. We further verify the efficacy of our approach by comparing with recently proposed inverse protein folding methods and by rigorous ablation studies.

## 1 Introduction

Computational protein design (CPD) broadly attempts two goals: (i) *inverse folding*, also known as *fixed-backbone design* which aims to produce novel amino acid sequences conforming to a predefined protein backbone structure and (ii) *de novo design* which seeks to develop sequences encoding proteins with some desired properties. Success in these areas has led to the development of enhanced therapeutics (Chevalier et al. 2017), biosensors (Quijano-Rubio et al. 2021), enzymes (Siegel et al. 2010) and more (Lucas and T. Kortemme 2020; Tinberg et al. 2013).

Owing to the conventional wisdom that a protein’s native state corresponds to its free energy minimum, CPD tasks are traditionally framed as an energy minimization problem. In this setting, the energy function typically consists of some combination of physics-based terms (J.H. et al. 2018; Alford et al. 2017; X. Huang et al. 2019; Junmei Wang et al. 2004) and knowledge-based terms (Park et al. 2016; C. Zhang et al. 2004; Zhou et al. 2020; Xiong et al. 2014), the latter of which are often derived from experimental data. During optimization, sequences are sampled and mutated until an energy minima is reached. Although this approach has garnered some success, it has a few major drawbacks. First, the size of the search space increases exponentially in the designed sequence length. This presents considerable challenges for designing large proteins. Next, for computational efficiency, score terms regularly approximate the total energy as the sum of weighted one and two-body terms. As a consequence, more complicated many-body interactions are ignored. Moreover, the extent to which designed sequences reflect their native analogs is limited by the accuracy of the underlying score function.

Over the past decade, computational biology has seen a spate in research replacing traditional physics and knowledge-based approaches with machine learning (ML) methods. In terms of structural modelling, the recent advent of rotation-invariant (Schütt et al. 2018; Andrearczyk et al. 2020) and rotation-equivariant architectures (Thomas et al. 2018; Fuchs et al. 2020; Satorras et al. 2021; Jing et al. 2021; Weiler et al. 2018) for learning from 3D-coordinates has helped facilitate explicit representations of geometric information, which has in turn led to improved performance in molecular modelling tasks. Most notably, AlphaFold2 (Jumper et al. 2021) and RoseTTAFold (Baek et al. 2021) introduced two-stage architectures, relying on equivariant neural networks and innovative attention mechanisms, which are capable of accurately predicting the structure of many novel proteins. Given the success of these architectures, coupled with the inherent geometric nature of inverse protein folding, we sought to develop a similar architecture for this task.

Parallel to the inverse folding task, we also sought to understand the extent to which protein function can be captured from structural information alone. As protein function is jointly determined by its 3-dimensional conformation and primary amino acid sequence (Mitchell et al. 2019; Dawson et al. 2017) this suggests that functional effects of mutations should be modeled jointly as well. It has been shown that protein structure information can be used to predict its function (Gligorijevic et al. 2021; Lai and Xu 2022). Although the most successful existing methods for protein mutation effect prediction rely on sequence and evolutionary information (Meier et al. 2021; Hopf et al. 2017; Riesselman et al. 2018), we hypothesized that a generative model, conditioned solely on structure and partial sequence, could be used as a zero-shot predictor for functional effects on single point mutations.

Here, we extend our recent work on side-chain coordinate prediction (McPartlon and Xu 2022) and introduce a deep SE(3)-equivariant graph transformer architecture which simultaneously predicts each residue’s identity and side-chain conformation. We compare to several existing inverse folding methods on CASP13, CASP14, CATH4.2, and TS50 test sets and show that our method achieves significantly higher native sequence recovery (NSR) rates across all datasets. In addition, we verify our models efficacy in capturing functional aspects of proteins by comparing predicted likelihoods against Deep Mutational Scanning (DMS) experiments. This result sheds light on the use of structure information in future studies on protein mutation effect prediction. To achieve these results, we explore several novel training and loss strategies and demonstrate their effectiveness through careful ablation studies.

## 2 Related Work

Several DL-based approaches have been proposed for inverse protein folding. These methods almost invariably employ convolutional neural networks (CNNs) or Graph Neural Networks (GNNs) as their underlying architecture. For a given architecture, the primary differences between approaches mainly stems from their feature representations; namely the representation of backbone geometry. We now give a brief overview CNN and GNN-based methods for inverse folding. For a more detailed review of feature representations applied to CPD models, see (Defresne et al. 2021).

Several CNN-based architectures have been applied to computational protein design (Yi Zhang et al. 2019; Qi and J. Z. H. Zhang 2020; Chen et al. 2020; Anishchenko et al. 2021; Shroff et al. 2019). In the fixed-backbone setting, Chen et al. employ 2D-CNNs in SPROF (Chen et al. 2020), which incorporates two-dimensional pairwise distance features to improve on the performance its predecessor SPIN2 (O’Connell et al. 2018). More recently, ProDCoNN (Yi Zhang et al. 2019), and DenseCPD (Qi and J. Z. H. Zhang 2020) applied convolutions to 3D volumetric regions of the input, allowing sequential features to be represented with rotation invariant voxel encodings of residue microenvironments. This approach has the benefit of allowing the network to extract spatial and geometric features without delegating to hand-crafted feature representations used in the 2D architectures.

In contrast to CNN architectures, GNN-based methods treat the input protein as an augmented graph, with features attached to each node (residue) and pair of nodes (edge). Some examples include the GNN variant of geometric vector perceptrons (GVP-GNN) (Jing et al. 2021), the Structure Transformer (Ingraham et al. 2019), ABACUS-R (Liu et al. 2022), and ProteinSolver (Strokach, Becerra, et al. 2020). These methods parallel language models, where each residue identity is iteratively predicted by conditioning on an encoded representation of the input graph. With the exception of ProteinSolver, each of these methods attempts to explicitly model 3D coordinates. The Structure Transformer, and Abacus-R accomplish this using relative orientation encodings of invariant local coordinate systems. On the other hand, the SE(3)-equivariance of GVP’s enables GVP-GNN to operate directly on backbone coordinates as input. To the best of the author’s knowledge, GVP-GNN is the first fully equivariant architecture applied to inverse protein folding.

Akin to fixed-backbone design, several DL-based architectures have been applied to de novo design (Jue Wang et al. 2021; Anishchenko et al. 2021; Lin et al. 2022; Jin et al. 2022). For general reviews and background on de novo and fixed-backbone design, we refer the reader to (Castorina et al. 2021; Ovchinnikov and P.-S. Huang 2021; Coluzza 2017; Pan and Tanja Kortemme 2021). For a more complete review of model classes applied to protein design, see (Strokach and Kim 2022).

Alongside CPD, considerable progress has been made in predicting the effect of point mutations on protein stability. These methods often rely on sequence evolutionary information such as EVmutation (Hopf et al. 2017), DeepSequence (Riesselman et al. 2018), SIFT (Ng and Henikoff 2003)), and PolyPhen-2 (Adzhubei et al. 2013) or large language models trained on massive sequence databases such as ESM-1v (Meier et al. 2021). While these methods have considerable merit, they inherently rely on adequate evolutionary information to make predictions. In contrast, our structure conditioned sequence model can be used as a zero-shot predictor for mutation effects when evolutionary information is absent.

Aside from CPD-specific models, our architecture draws from recent advancements in DL-based methods for 3D-modelling and protein structure prediction. Notably, we utilize memory efficient variants of triangle multiplication and triangle attention introduced in AlphaFold2 along with a TFN-based SE(3)-equivariant transformer inspired by the SE(3)-Transformer of Fuchs et al (Fuchs et al. 2020). Our work on side-chain coordinate prediction (McPartlon and Xu 2022) details these modifications, which were used to construct a deep GNN for protein side-chain packing.

## 3 Methods

In this work, we model the inverse folding problem with an SE(3)-equivariant architecture operating directly on features derived from backbone coordinates. Our architecture consists of two main sub-modules. The first is a deep Locality-Aware Graph Transformer which utilizes the geometry of the input backbone to refine node and pair features and restrict attention to spatially local residue pairs. The output of this component, along with the input backbone coordinates, are passed to a deep TFN-transformer which produces side chain conformation and sequence identity for each input residue. The details of these architectural component are fully described in (McPartlon and Xu 2022), and a schematic overview is given in Figure 1.

**Figure 1:**
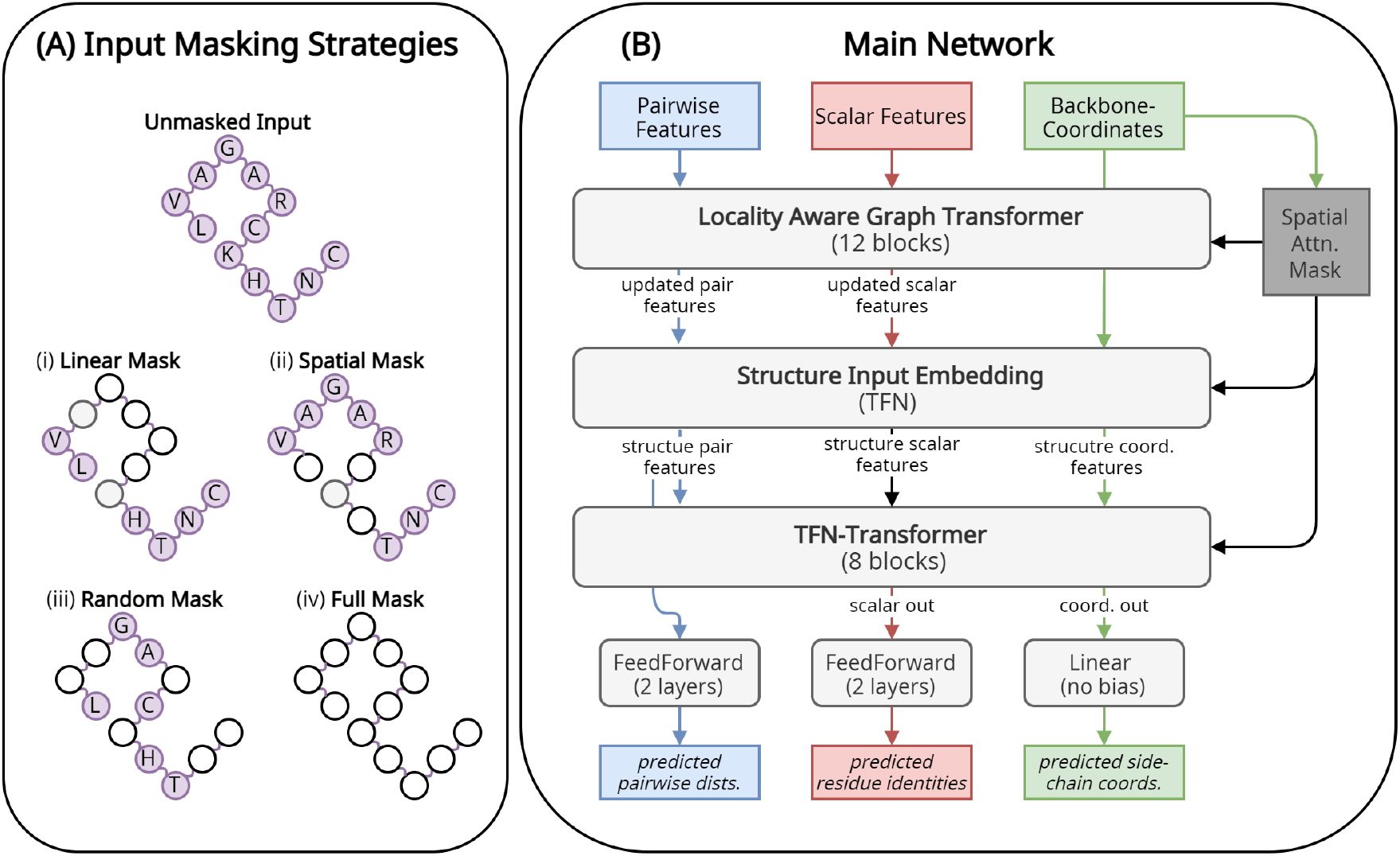
Approach Overview. **(A)** Our method employs several different strategies for masking input residue identities. Our best performing model employs one of (*i-iv*), sampled uniformly at random during training. Further details of each strategy can be found in Section 3.2. **(B)** Our deepest model consists of two main components: A 12-layer Locality Aware Graph Transformer, and 8-layer TFN-Transformer. Aside from predicting residue identities, the network also predicts side-chain conformations and pairwise distances between 6 pairs of atom types (described in Section S1.4).

### 3.1 Input Representation

We represent a protein as a graph 𝒢= (𝒱, *ε*) where each node *v*_*i*_ ∈ 𝒱 represents a residue, and an edge {*e*_*ij*_}_*i*≠*j*_ ∈ *ε* exists between each residue *i* and it’s *k* nearest neighbors computed by pairwise *Cα* distance. The number of nearest neighbors differs for the two components of our network. We shoose *k* = 30 for our first submodule, and *k* = 16 for the TFN-Transformer.

A brief description of scalar, coordinate, and pairwise input features is given below. We note that all input features are derived directly from backbone coordinates. Our input embedding procedure, along with a more detailed description of input features can be found in Figure S2, and Table S2.

### Residue Features

Scalar residue features are comprised of sin and cosine encodings of backbone dihedral angles, and one-hot encodings of each residue’s relative sequence position, and a one-hot encodings of residue identity (when applicable). A separate <mask> token is used for masked residues whose identities are left to be determined.

### Pair Features

Input pair features consist of one-hot encodings of one-hot encodings for pairwise distance between atom pairs (*Cα, Cα*), (*Cα, N*),, sin and cosine encodings of dihedral and planar angles defined by Yang et al (Yang et al. 2020), and a joint embedding of signed relative sequence separation *i*− *j* and pairwise residue-types (when applicable). To produce pairwise orientation information, we impute a unit vector in the direction *Cβ*_*i*_ − *Cα*_*i*_ before computing the respective angles. The imputed vector is calculated as in (Jing et al. 2021) using

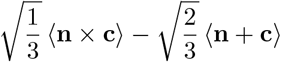

where ⟨*x*⟩ = *x/*∥*x*∥_2_, **n** = *N*_*i*_ − *Cα*_*i*_, and **c** = *C*_*i*_ − *Cα*_*i*_.

### Coordinate Features

For a given backbone conformation *X*, let *X*_*i*_ denote the coordinate of atom *X* in residue *i*. Input coordinate features for residue *i* consist of unit vectors *N*_*i*_ − *Cα*_*i*_, *C*_*i*_ − *Cα*_*i*_, and *O*_*i*_ − *Cα*_*i*_. This input is passed through a TFN to produce a hidden encoding of dimension 16. Note that only the TFN-Transformer component of our architecture operates on coordinate features. Furthermore, we use the raw positions of backbone *Cα* coordinates to compute an initial (equivariant) basis for our TFN-Transformer module.

### 3.2 Masking Strategies

While the ultimate goal of inverse folding is to predict sequence given structure alone, we hypothesized that providing the model with partial sequence information during training could lead to improved results. This decision was motivated by Ingraham et al., who demonstrate that unconditional language models struggle to assign high likelihoods to sequences from out-of-training folds (Ingraham et al. 2019). On the other hand, the authors show that conditional autoregressive modelling helps facilitate adaptation to specific and potentially novel parts of structure space. In light of this, we developed several masking strategies, allowing our model to condition on partial sequence information via input scalar and pair features during training (see Figure S2). The strategies are described below. We use *L* to denote sequence length, and use the notation *x* ~_*R*_ 𝒮 to denote uniform random sampling from the set 𝒮.

#### Spatial

A random residue *i* ~ _*R*_ [1..*L*] is chosen, and the identity of this residue, along with the *k* nearest neighbors of this residue are masked. Nearest neighbors are computed by pairwise *Cα* distance. In practice, we chose *k* = *pL*, where *p* ~_*R*_ (0.1, 0.5).

#### Linear

We first choose a length *𝓁* ~_*R*_ [*m*..*M*] and then choosing an index *i* ~_*R*_ [1..*L* − *𝓁*]. We then mask all residue identities in the range *i*..*i* + *𝓁* − 1. In choosing the length *𝓁*, we set *m* = 0.25*L*, and *M* = 0.75*L*.

#### Random

A threshold probability *p* ~ _*R*_ (0, 1), is sampled, and then each residue’s sequence identity is masked independently with probability *p*.

#### Full

All residue sequence identities are masked, and no sequence information is given to the model. This is the typical strategy used for inverse protein folding.

To evaluate the effect of masking strategy on model performance, we train four separate models using each strategy (independently), and an additional model which chooses one of the four strategies with equal probability for each training sample. Detailed results are given in Section 5.1.

### 3.3 Zero-shot Protein Mutation Effect Prediction

To quantify our models ability to predict protein mutation effects, we use the log ratios of the wild-type amino acid and the mutated amino acid at the mutated index *i*. Formally, for a protein sequence *S* = *s*_1_, … *s*_*n*_ and backbone conformation *C*, we compare the log probability ratio of mutation *x*_*mutant*_ appearing at sequence position *i* against the wild-type *x*_*wild−type*_ while conditioning on the backbone conformation *C*. and *S*_*−i*_; the identities of all amino acids aside form *s*_*i*_. The calculation is shown in Equation (1).

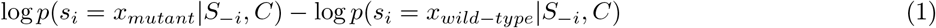

### 3.4 Architecture and Training Details

We briefly overview our model architecture, loss, and training details. Unless otherwise stated, all of our models were trained and validated using chains from the publicly available BC40 dataset. Our training set contains roughly 37k chains filtered to 40% nonredundancy. A more complete overview along with details of train and test set similarity, optimization procedure, hyperparameters, and input embedding, can be found in Section S1.

We trained two main variants of our model which we refer to as *small* and *large*. For each variant, self-attention and triangle updates are restricted to the same number of spatially-nearest neighbors. Our small architecture uses a hidden dimension of 120 for residue and pair features, and a hidden dimension of 16 for coordinate features. This variant consists of eight Locality-Aware Graph Transformer blocks stacked on six TFN-transformer blocks. The large model uses the same dimensions for pair and coordinate features, but increases the hidden dimension of residue features to 180. The submodule depths are increase to twelve and eight layers respectively.

The same training and loss strategies are employed for each variant. we randomly select one of the masking strategies overviewed in Section 3.2 for each sample, and compute three separate loss terms, one for each output feature, corresponding to predicted residue identity, pairwise distance between side-chain atoms, and predicted side-chain conformation. The latter two terms are given weight 0.15 relative to the predicted identity loss term. More details on the loss function can be found in Section S1.4.

## 4 Results

We begin by comparing our method to GVP-GNN (Jing et al. 2021), DenseCPD (Qi and J. Z. H. Zhang 2020), ProDCoNN (Yi Zhang et al. 2019), ProteinSolver (Strokach, Becerra, et al. 2020), and Rosetta Design (Chaudhury et al. 2010; J. K. Leman et al. 2020). Details on data collection for these five methods can be found in Section S2. Computational protein design methods are notoriously difficult to benchmark, as some sequences may recapitulate nearly identical structures, while some structures may have no conforming sequences. In line with Ingraham et al., we report (i) native sequence recovery (NSR) rate, and (ii) model perplexity (when applicable). The rationale is that NSR assesses how closely a designed sequence matches the native sequence of an input backbone structure, and perplexity tests the ability of the model to assign a high likelihood to the native sequence. Following Jing et al. (Jing et al. 2021), NSR is reported as the median (over all structures) of the average percentage of residues correctly recovered.

### 4.1 CASP13 and CASP14 Targets

#### Recovery and Perplexity

We chose to evaluate each method on CASP13 and CASP14 test sets (see Table S3 for full list of targets) as there is no canonical training or validation sets for fixed-backbone design, and in most cases, these test sets have little overlap with the training sets used for the methods in comparison. We empirically verify this claim by comparing sequence and structural similarity of all training chains in CATH4.2 (curated by Ingraham et al. (Ingraham et al. 2019)) and BC40 against target chains for each test set in Section S2.1. Although no standardized test sets exist, CATH4.2 and TS50 have emerged as de-facto benchmarks used by several methods. In light of this, we include a comparison in our extended results (Figure S6, and Figure S5) (see Table S3 for a list of these targets).

Our method outperforms all competitors in terms of NSR accross both datasets. Interestingly, DenseCPD retains the second highest recovery scores on the CASP14 dataset while having much higher perplexity than our method and GVP-GNN. To better understand this, we further compare our method with GVP-GNN and DenseCPD in terms of precision, recall, and F1-score for each amino acid type on these datasets.

#### Precision, Recall, and F1

From Figure 2 we see that our method achieves top F1 scores for the majority of amino acids (13/20). DenseCPD also performs well, achieving top F1-scores for the remaining seven residue types. Not surprisingly, all three methods perform well in predicting glycine (G) and proline (P) which can be attributed to the unique structural features of the two amino acids.

**Figure 2:**
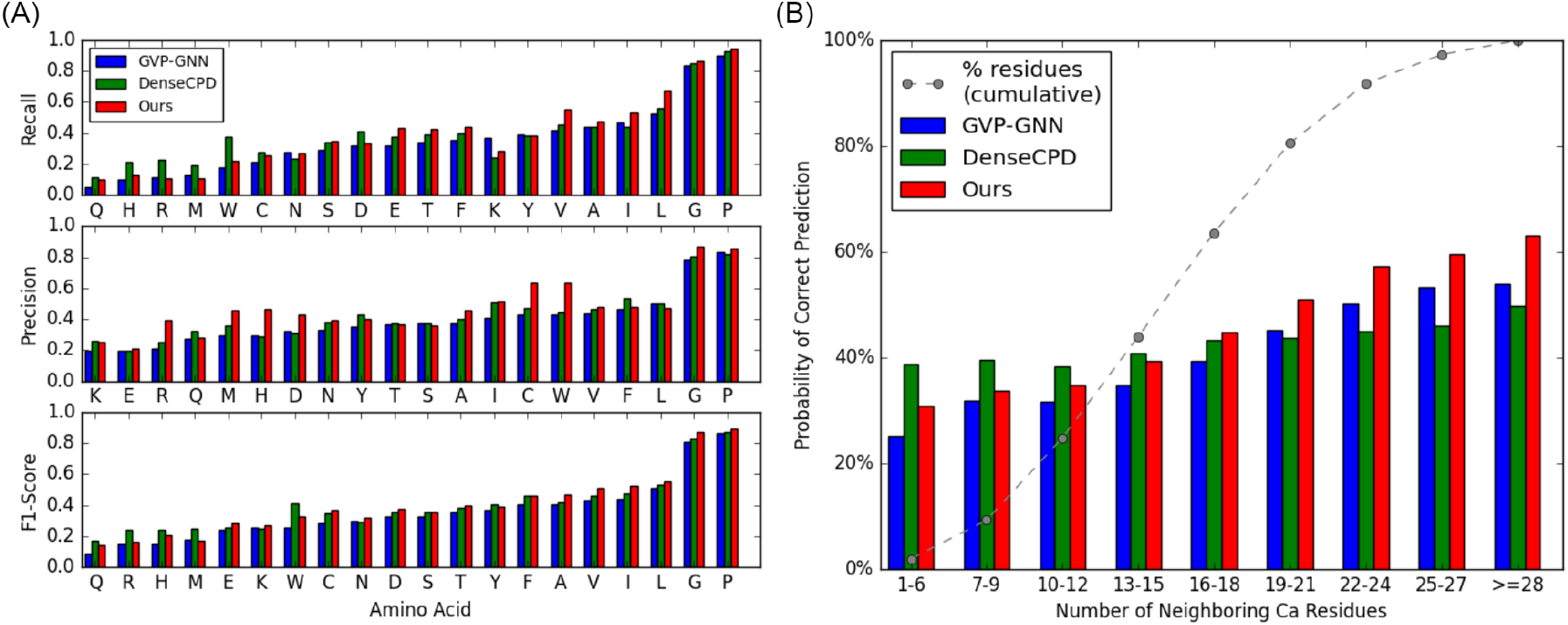
Comparison of GVP-GNN, DenseCPD and Our method on CASP13 and CASP14 targets. (**A**) Precision, Recall and F1-score for all 20 amino acid types; F1-score is computed as 2*pr/* (*p* + *r*) where *p* and *r* denote precision and recall respectively. (**B**) Comparison of prediction accuracy conditioned on number of residues in the target’s immediate microenvironment. The x-axis shows the number of residues whose *Cα* atom is within 10Å of the target residue. The dotted-gray line shows the cumulative percentage of residues in the dataset with (up to) the number of neighbors listed on the x-axis.

In terms recall, we see that DenseCPD outperforms GVP-GNN for most residue types (16/20), and exceeds our method for 7/20 residue types. In contrast, we surpass DenseCPD in terms of precision for 5/7 residue types where DenseCPD achieves higher recall. As precision measures the number of correct predictions normalized by the number of predictions of a given type, and recall measures the proportion of correct predictions, this analysis aids in understanding why DenseCPD achieves high perplexity while maintaining high prediction accuracy.

We also compare the accuracy of the three methods conditioned on residue degree centrality; the number of neighboring *Cα* atoms in the target amino acid’s immediate microenvironment. This measure correlates with solvent accessible surface area, and acts as a proxy indicator of whether the corresponding amino acid is closer to the protein surface (low centrality) or core (high centrality). Analogous to Zhang et al. (X. Huang et al. 2020) we define core residues those amino acids with at least 20 *Cα* atoms within a 10Å radius of the target residue’s *Cα* atom. Similarly, surface residues are defined as those amino acids with as most 15 *Cα* atoms within the same region.

When conditioned on centrality, DenseCPD outperforms our method and GVP-GNN for surface residues residues, but quickly falls behind for residues closer to the protein core. For surface and core residues, GVP-GNN and our method improve their accuracy from 33% to 48% and 36% to 55% respectively. On the other hand, DenseCPD does not obtain large accuracy improvements, improving from 40% to 45% when considering the same criteria. This observation may stem from the difference in modelling approaches used by the two methods. Whereas our method and GVP-GNN model the input as a graph and jointly update residue features via self-attention or message passing, DenseCPD models joint interactions only at the input level; updating hidden features independently for each residue.

### 4.2 Zero-shot mutation effect prediction

To better understand the extent to which our model’s designs capture functional aspects of the underlying protein, we compare predicted log likelihoods assigned to point mutations using stability data gathered from several DMS datasets.

Following Ingraham et al. (Ingraham et al. 2019), we first assess our models performance on predicting protein stability in response to single point mutations using deep mutational scanning data from (Rocklin et al. 2017). In Table 2 we observe that conditioning on the partial input sequence (in addition to backbone conformation) leads to improvements in stability prediction for eight of the ten de novo designed mini-proteins. Moreover, our model improves over the Structure Transformer for nine of the ten targets.

**Table 1:** Results for CASP13 and CASP14 targets. We report median NSR (higher is better) and median perplexity (lower is better) over target chains in CASP13 and CASP14 datasets. Perplexities are omitted for ProteinSolver and Rosetta. The former calculates log probabilities only for the predicted residue type, and the latter is energy-based. Methods GVP + BC40 and GVP + CATH4.2 use the same hyperparameters as reported in (Jing et al. 2021), but differ on the training set used - CATH4.2, and BC40, respectively.

|  | CASP13 |  | CASP14 |  |
| --- | --- | --- | --- | --- |
| | Recovery ↑ | Perplexity ↓ | Recovery ↑ | Perplexity ↓ |
| Ours (Large) | **50.6%** | <u>4.88</u> | <u>41.1%</u> | <u>6.17</u> |
| Ours (Small) | <u>48.9%</u> | **4.82** | **41.3%** | <u>6.17</u> |
| DenseCPDNet<sup>1</sup> | 48.1% | 5.14 | 39.0% | 7.18 |
| GVP + BC40 | 43.3% | 5.04 | 38.3% | **5.58** |
| GVP + CATH4.2 | 44.0% | 5.12 | 36.4% | 6.40 |
| ProteinSolver | 37.0% | - | 32.8% | - |
| SPROF | 37.4% | 7.30 | 33.8% | 8.02 |
| Rosetta | 34.2% | - | 27.1% | - |

**Table 2:**
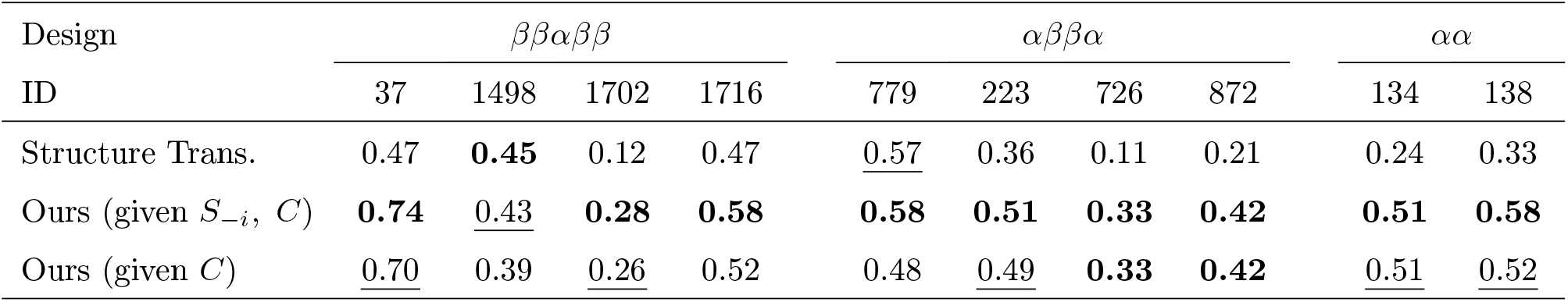
Predicted likelihoods correlate with mutation effects in de novo - designed mini proteins. Pearson Correlation between high-throughput mutation effect data from a systematic design of miniproteins. Each design (column) includes 775 experimentally tested mutant protein sequences. Results are split by fold topology (first row) and design model (second row) as referenced in (Rocklin et al. 2017). Results for the Structure Transformer are generated on rigid backbones, and taken from Table 5 of (Ingraham et al. 2019).

| Design | $\beta\beta\alpha\beta\beta$ | | | | $\alpha\beta\beta\alpha$ | | | | $\alpha\alpha$ | |
| --- | --- | --- | --- | --- | --- | --- | --- | --- | --- | --- |
| ID | 37 | 1498 | 1702 | 1716 | 779 | 223 | 726 | 872 | 134 | 138 |
| Structure Trans. | 0.47 | **0.45** | 0.12 | 0.47 | <u>0.57</u> | 0.36 | 0.11 | 0.21 | 0.24 | 0.33 |
| Ours (given $S_{-i}$, $C$) | **0.74** | <u>0.43</u> | **0.28** | **0.58** | **0.58** | **0.51** | **0.33** | **0.42** | **0.51** | **0.58** |
| Ours (given $C$) | <u>0.70</u> | 0.39 | <u>0.26</u> | 0.52 | 0.48 | <u>0.49</u> | **0.33** | **0.42** | <u>0.51</u> | <u>0.52</u> |

Moreover, our model outperforms protein language models TAPE (Rao et al. 2019) UniRep (Alley et al. 2019) trained on large sequence databases across 12 DMS datasets in zero-shot protein mutation effect prediction regardless of sequence evolutionary information as shown in Figure 3(A). This result indicates that our method captures the underlying functional interaction between 3-dimensional conformation and amino acid sequences and suggests structure information may facilitate the characterization of protein mutation effects. We also noticed in Figure 3(B) that the performance of our method correlates with the similarity between the DMS target structure and our training set but independent of the sequence identity between the DMS target and our training set. This further confirms that our zero-shot mutation effect prediction is structure-aware.

**Figure 3:**
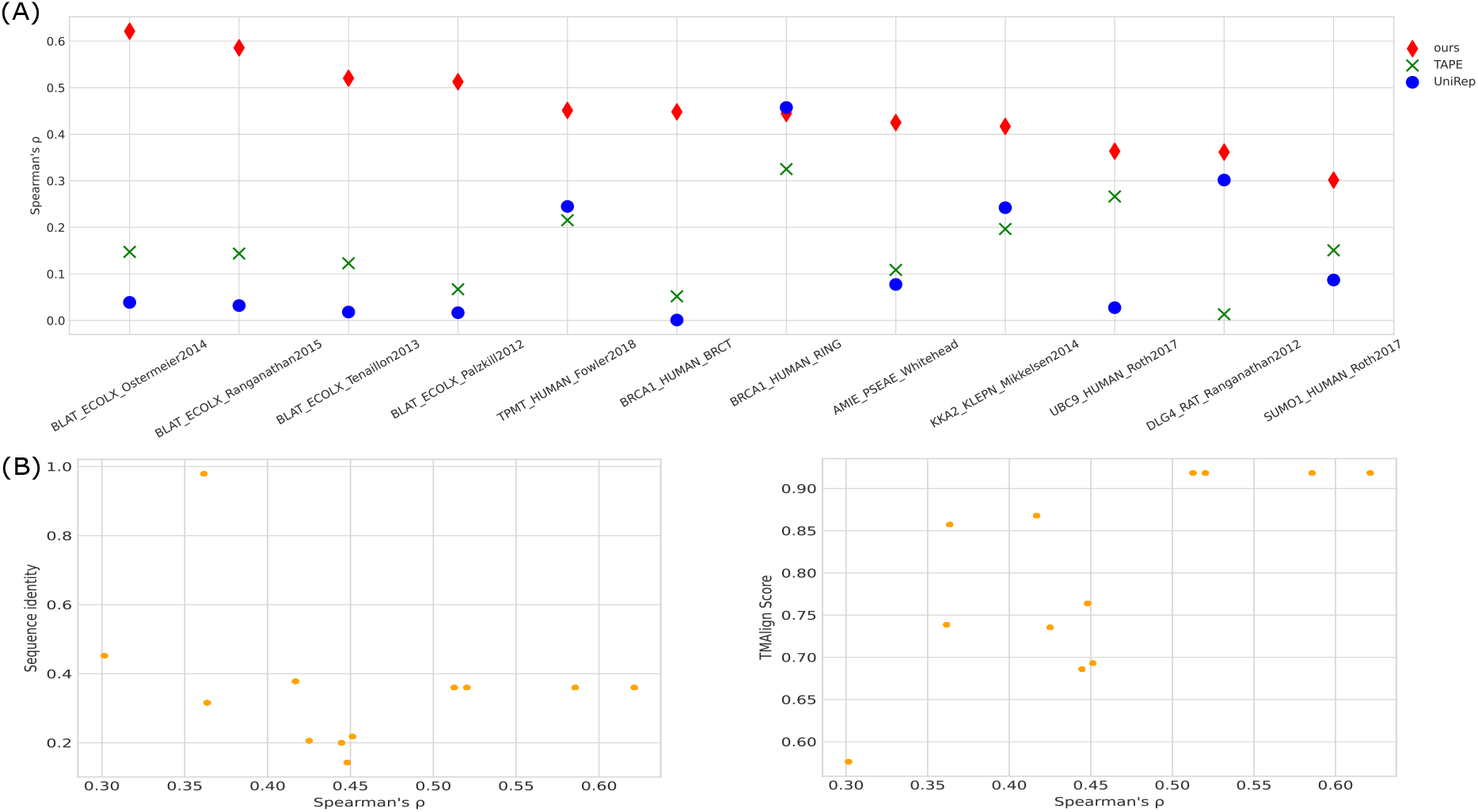
Zero-shot prediction performance. (**A**) Per task performance of our model compare to two other language models UniRep and TAPE on 12 DMS datasets measured by rank correlation. (**B**) Performance of our model against the top-1 TMAlign score(top) and sequence identity(bottom) among the training protein and DMS targets.

## 5 Ablation Studies

To validate the impact of our training strategy, and to understand the influence of particular architectural components, we conducted ablation studies focused on our masking techniques, loss, two-stage architecture, and model hyperparameters.

### 5.1 Mask Strategy Evaluation

In Section 3.2 we outlined four different masking strategies. We now evaluate the performance of five different models *Spatial, Linear, Random, Full*, and *All*, each of which was trained using only input derived from the respective strategy (all strategies were used for *All)*.

In Table 3 we see that both recovery (NSR) and perplexity are improved by training with an equal combination of our four masking strategies. Most notably, the perplexity of the model trained with all mask types is significantly lower than those trained with a single mask type across all test strategies. When evaluating on fully-masked input, the corresponding training strategy produces second-best results for both NSR and perplexity. Of the other three single-mask input strategies, the random masking strategy performs best overall for the testing criteria. Surprisingly, the random strategy achieves similar results or outperforms spatial and linear masking on the corresponding test input.

**Table 3:** Comparison of Masking Strategies on CASP13 targets. The *Training Strategy* (rows) refers to the masking type used to train each model. The *Test Strategy* (columns) refers to the type of mask applied to the input to obtain the results. The cell values indicate the median NSR rate and perplexity for the corresponding mask-type pair. For a given test strategy, ten separate masks were applied to each test chain, and the recovery/NSR values for the chain was taken as the average over the ten samples (with the exception of full-masking). The same masks were used to evaluate each training strategy. For all results, recovery (NSR) and perplexity scores are evaluated only on the masked region of the input chain. We performed a paired t-test to confirm that the combined masking strategy (All) significantly outperformed the other four models in terms of NSR and perplexity and obtained p-values less than 0.001 for all tests.

|  |  | Test Strategy |  |  |  |  |  |  |  |
| --- | --- | --- | --- | --- | --- | --- | --- | --- | --- |
| | | NSR ↑ | | | | Perplexity ↓ | | | |
|  |  | Spatial | Linear | Random | Full | Spatial | Linear | Random | Full |
| **Train**<br>**Strat-**<br>**egy** | Spatial | 45.6% | <u>46.3%</u> | 46.3% | 43.5% | 5.26 | 5.10 | 5.12 | 5.80 |
|  | Linear | 46.4% | 46.1% | 45.0% | 43.5% | 5.25 | 5.05 | 5.25 | 5.69 |
|  | Random | <u>48.1%</u> | 46.2% | <u>47.0%</u> | 44.5% | <u>5.13</u> | <u>4.91</u> | <u>5.02</u> | 5.47 |
|  | Full | - | - | - | <u>46.6%</u> | - | - | - | <u>5.14</u> |
|  | All | **53.2%** | **49.9%** | **52.0%** | **48.9%** | **4.64** | **4.52** | **4.54** | **4.82** |

We also consider the impact of architectural choices in terms of model loss, hidden dimension of residue features, the use of transformer submodules, and model depth. In Figure 4(A), we see that removing side-chain RMSD (*vii*) and predicted pairwise distance between side-chain atoms (*viii*) from the loss function significantly degrades NSR for CASP13 targets. With the default loss held constant, removing the TFN-Transformer has the largest impact on NSR, which drops nearly five percentage points when this component is ablated. Surprisingly, our Locality-Aware Graph Transformer module (*Tri*) still slightly outperforms GVP-GNN for CASP13 targets (44.3% NSR vs. 44.0% NSR).

**Figure 4:**
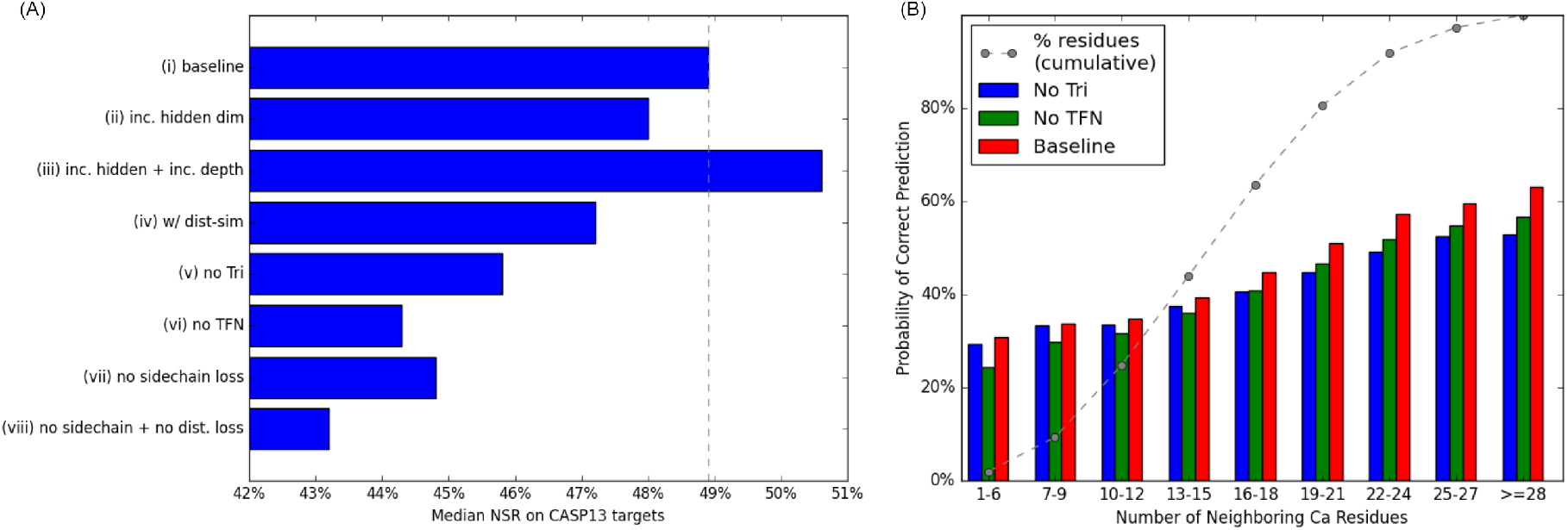
Impact of architectural components on NSR for CASP13 targets. (**A**) Contribution of features towards network performance; all ablations are taken with respect to the baseline small model model (*i*). The bars show the median NSR of the corresponding model on CASP13 targets. The gray dotted line shows the median NSR of *(i)*. (*ii*) shows performance when increasing the residue hidden dimension from 120 to 180 and (*iii*) when also increasing the depth from 14 to 20. (*iv*) shows the result of using distance vs. dot-product based coordinate attention. *(vi)* and *(vi)* remove the Locality-Aware Graph Transformer and the TFN-Transformer submodules, respectively. *vii* ablates side-chain coordinate loss and *viii* removes both side-chain and pairwise distance loss. (**B**) Performance of the baseline model (*i*) and models (*v*), (*vi*) where Triangle Updates or the TFN-Transformer are ablated (respectively). The gray dotted shows the cumulative percentage of residues with respect to the number of *C*_*α*_ neighbors (x-axis) in a 10Å radius of the central residue being predicted.

We further compare the benefits of our two-stage architecture in Figure 4(B). From this, we observe that the two stage architecture significantly outperforms each single component when conditioned on all nine levels of degree centrality. These results suggest that the representations learned by each submodule are at least partially disjoint - and there is a clear benefit to combining the two representations.

## 6 Conclusion

In this work, we developed and evaluated a framework for learning inverse protein folding. Our model uses features derived solely from protein backbone coordinates and further leverages this information to restrict attention to spatially-local residue pairs.Although our model does not make explicit use of evolutionary information, it is able to capture functional aspects of the underlying protein demonstrated by correlation with stability data gathered from deep mutational scanning studies and outperforms protein language models that trained on large sequence databases. We hope our model can prompt other researchers to further integrate structure information for characterizing protein mutation effects. On top of the model itself, we showed that our loss function and training strategy offer performance enhancements when compared to traditional approaches. The former effectively exploits the properties of SE(3)-equivariance and the latter facilitates generalization to novel parts of structure space.

## 7 Author Contributions

J.X. conceived and supervised the project, built the in-house training data and revised the manuscript. M.M. developed and tested the algorithm, analyzed and collected the results, and wrote the manuscript. B.L. conducted and analyzed the deep mutational scanning study and assisted in writing the manuscript.

## 8 Conflict of interest

The authors declare that they have no conflict of interest.

## 9 Code Availability

Code will be made publicly available upon formal publication.

## Supplementary Information

### S1 Training Details and Hyperparameter Overview

#### S1.1 Training

We trained and validated all models using the BC40 training dataset and validated on BC40 validation set (January 2020 release). The training and validation splits contain 37k, and 1.5k and chains (resp.), which are selected from PDB database by 40% sequence identity cutoff. In addition, we also filtered our training and validation sets against CASP13 targets, removing any chain with >40% sequence similarity as reported by MMSeqs (Mirdita et al. 2021). There is some discrepancy between sequence identity returned by MMseq and those computed by TM-Align (Y. Zhang and Skolnick 2005) shown in Figure S3 which is likely due to the (potentially) local alignment used by TM-Align.

All of our models were trained using early stopping based on validation loss, and for a total of at most 10 epochs. Because of the memory overhead of TFNs and Triangle updates, we used a sequence crop size of 300 to avoid running out of memory. Overall, each model was trained for roughly five days on a single nvidia RTX A6000 gpu.

#### S1.2 Input and Hyperparameters

In tuning our model, we mainly experimented with feature hidden dimensions and model depth. In the main text, we present results for *large* and *small* variants of our model. The parameter values for each are shown in Table S1.

**Table S1:** Hyperparameters. Model depth, and hidden dimensions for residue (*scalar*), coordinate, and pair features used in the Locality-Aware Graph Transformer and TFN-Transformer submodules of our network. items are listed separartely for small and large variants of our model when the values differ.

|  | Graph<br>Transformer<br>(*residue, pair*) | TFN-Transformer<br>(*scalar, coord.*) |
| --- | --- | --- |
| Depth | Large: 12<br>Small: 8 | Large: 8<br>Small: 6 |
| Hidden Dim. | Large : 180, 120<br>Small: 120, 120 | Large: 180, 20<br>Small: 120, 16 |
| Num. Attention<br>Heads | 10, 4 | 10, 10 |
| Head Dim. | Large: 20, 32<br>Small: 20, 28 | 20, 4 |
| Neighbor Distance<br>Cutoff | 15, 15 | 15 |
| Max. Nearest<br>Neighbors | 30, 30 | 16 |

#### S1.3 Input Features

In Table S2 we briefly review our model’s input features and their respective shapes. We note that each feature (aside from residue-type) can be derived entirely for protein backbone coordinates.

**Table S2:** Input features and shapes. Feature names and shapes are shown in the left column, and a description of each feature (row) is given in the right column.

| Features Name & Shape | Description |
| --- | --- |
| res_type<br>[L] | A number in the range 0..21 representing the corresponding amino acid type. The first 20 labels correspond to the 20 standard amino acid types, and the final label corresponds to a "missing" identity. |
| bb_dihedral<br>[L, 3] | Backbone phi, psi, and omega dihedral angles. |
| seq_pos<br>[L] | The sequence index of the respective residue, from 1..L |
| centrality<br>[L] | The number of $C\alpha$ atoms within 16Å of residue of the respective residue's $C\alpha$ atom. |
| atom_distance<br>[L, L, 3] | One-hot encoded binned distance between $C\alpha - C\alpha$, $C\alpha - N$, and $N - O$ atoms in each residue. Each bin represents a distances from 2Å-20Å with two separate bins used for distances falling outside of this range. |
| tr_orientations<br>[L, L, 3] | Dihedral and planar angles defined by Yang et al (Yang et al. 2020). |

Our input features are embedded following the strategy proposed in (McPartlon and Xu 2022). The primary difference is that we we remove one-hot encodings of angle-based features. add an extra token to the residue type input representing a “masked” residue identity. The full procedure can be found in Figure S2. We note that the extra “masked” residue token enables us to generate designs in the classical setting, from backbone information alone.

Figure S1: Input Embedding

**Figure S2:**
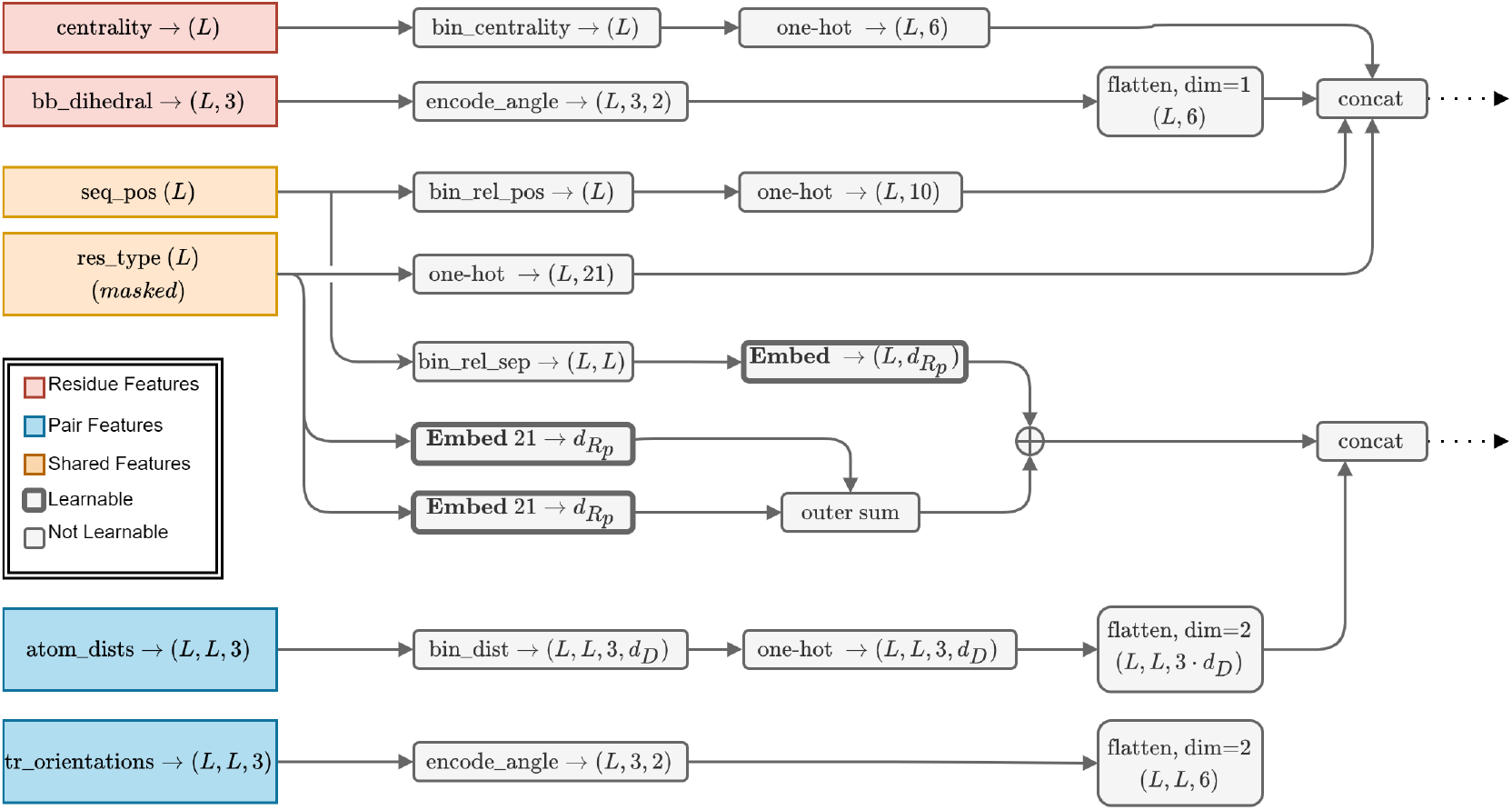
Input Embedding. The input feature embedding concatenates and projects residue and pair features directly to the hidden dimensions specified in Table S1. For one-hot encodings of distance, we use an encoding dimension of *d*_*D*_ = 32 for all models in the manuscript. For relative sequence separation and residue type encodings we use an encoding dimension of 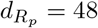

#### S1.4 Loss

Three separate loss functions are applied to the output - one for each of the predicted residue, pair, and coordinate features. Cross entropy (NSR) loss is computed on predicted residue features after applying a shallow feed-forward network to produce residue-type logits. Notably, we compute loss on the predicted residue feature even if the corresponding amino acid type was included in the input. Pair features are treated similarly, the output is passed through a shallow feed-forward network to obtain predicted pairwise distance logits between each *Cα* and side chain atoms *Cβ, Cγ, Cγ*_1_ *Cγ*_2_ *Oγ*_1_. Masked cross entropy loss is applied to this output based on the corresponding native structure’s residue type. Side chain conformation loss is computed as in (McPartlon and Xu 2022). The pair and side-chain conformation loss are given a weight of 0.15, and the NSR loss is given a weight of 1.

#### S1.5 Optimization

To optimize our models, we use Adam (Kingma and Ba 2015) with and an initial learning rate of 10^−3^and parameters *β*_1_ = 0.9, *β*_2_ = 0.999, and *ϵ* = 10^−8^, and use a minibatch size of 24. To stabilize training we apply gradient clipping by global norm to clip the gradients of each example in a minibatch to have *𝓁*_2_ norm at most 1. For every residual connection in our transformer blocks we use ReZero (Bachlechner et al. 2020), initialized with *α*_0_ = 0.1. To conserve GPU memory, we also apply gradient checkpointing on the triangle attention logits and TFN kernel outputs.

### S2 Data Collection

Results for ProteinSolver were generated following the instructions on the author’s github page. We ran the program with pre-trained model 191f05de/e53-s1952148-d93703104.state, using the *A*^*∗*^ search strategy until 1k sequences were designed, or 10k iterations of search were performed (whichever came first). The designed sequence with the smallest cumulative log-probability was chosen as the design for comparison. Results for GVP-GNN-CATH4.2 were generated using the pretrained model available on the author’s github page (here). Results for GVP-GNN-BC40 were gathered by retraining the model using the training loop from the same github repo with the default hyperparameters described in (Jing et al. 2021). Results for DenseCPD were collected by correspondence with the authors. All results for RosettaDesign were generated using Pyrosetta4, running fixbb protocol with flags *-ex1 -ex2 -ex3 -ex4 -multi_cool_annealer 10 -minimize_sidechains -linmem_ig 1 -nstruct 10* using the ref2015 score function. With these settings, ten designs are generated for each target, and we choose the design with the lowest energy for comparison.

#### S2.1 Train and Test Similarity

No canonical training or test dataset exists for fixed backbone design. Still, recent ML-based fixed-backbone design methods have been trained on CATH4.2 dataset. In light of this, we provide an assessment of sequence and structural similarity (see Figure S3) between CATH4.2 and BC40 datasets to the CASP13 and CASP14 test sets used in our results. Overall, the chains in each of the two training sets have comparable sequence and structural similarity with the most top-1 most similar CASP target chains.

**Figure S3:**
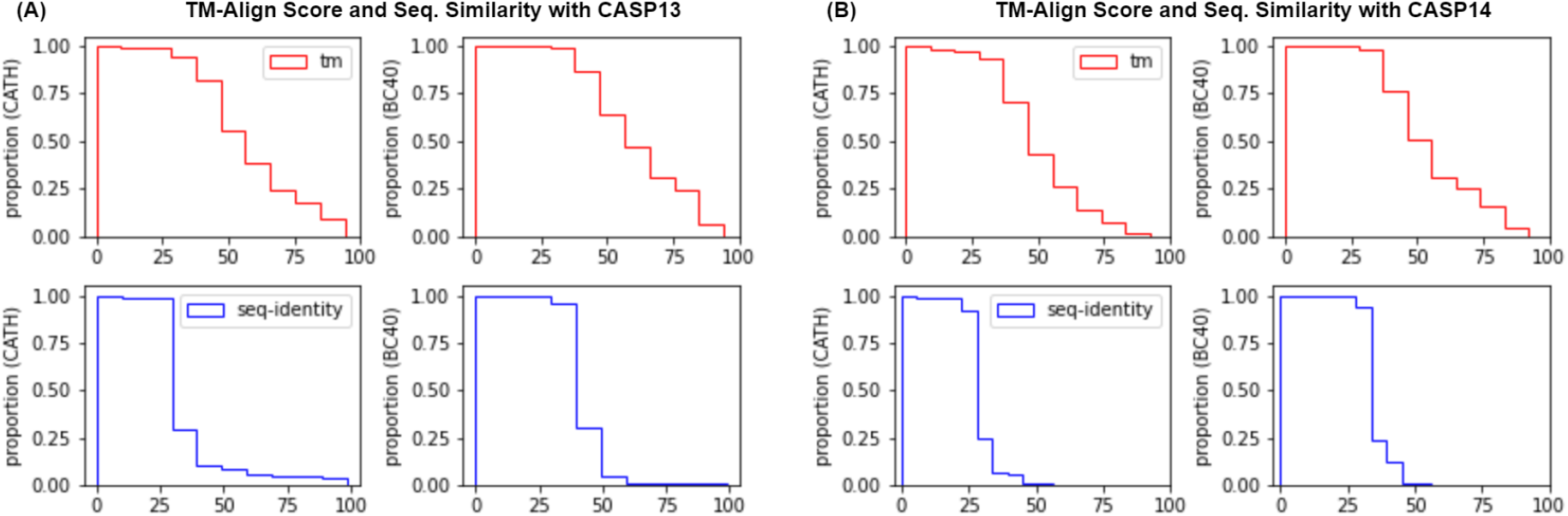
Cumulative frequency plot of top-1 TM-Align score and sequence identity from CASP13 and CASP14 test sets to CATH4.2 and BC40 training sets. For each plot, the x-axis shows similarity - TM-score (red) or sequence identity (blue) - of test targets to the most similar training target in CATH4.2 or BC40 datasets. The y-axis shows the cumulative proportion of similar targets. For example, the upper left plot tells us that roughly 20% of targets in CASP13 have TM-score greater than 75% to some target in CATH4.2. TM-scores and sequence identity were computed using TM-Align. For each target in the test sets, the sequence identity and TM-Align score of the most-similar model in the training set was added to the frequency plot.

We repeated this procedure for the 10 chains in our DMS test dataset used to evalute zero-shot mutation effect prediction. The results are show in Figure S4.

**Figure S4:**
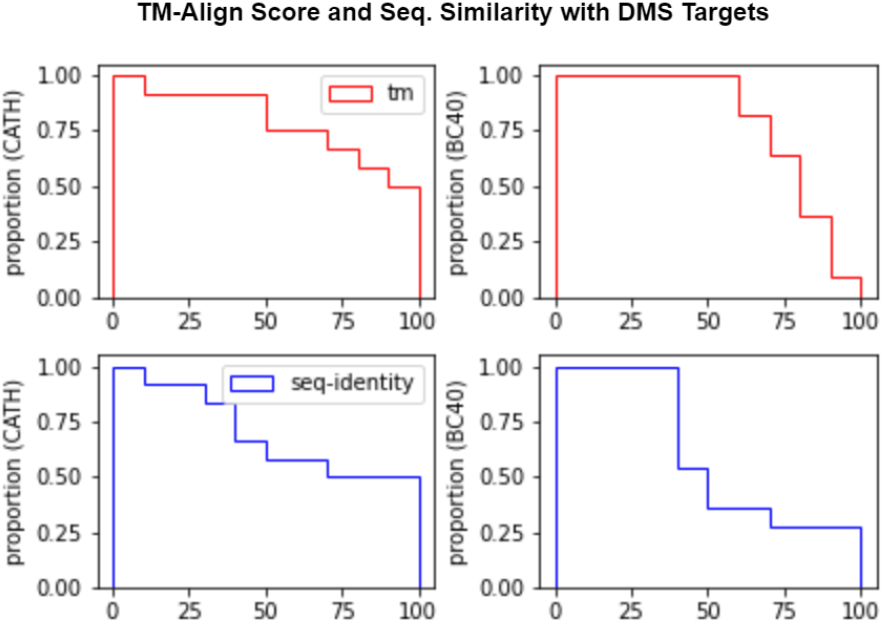
Cumulative frequency plot of top-1 TM-score and sequence identity from our DMS test dataset to CATH4.2 and BC40 training sets. The same procedure as performed for Figure S3 was repeated for the ten targets in our DMS test dataset.

We remark that, although our model out-performs GVP-GNN on the zero-shot mutation effect prediction task, the CATH4.2 training set has larger overlap with these targets in terms of both structural and sequence similarity.

### S3 Extended Results

Although there is no canonical test set for fixed-backbone design, TS50 and CATH4.2 have emerged as de-facto benchmarks for recent ML-based methods. We compare our model on the TS50 test set with several other ML-based design methods and physics-based Rosetta fix_bb application using results reported from the respective manuscripts. To obtain our results, we filter our BC40 training set to exclude all sequences with global similarity greater than 40% as computed with MMseqs (denoted with “filtered”).

#### Results for TS50 Test Set

**Figure S5:**
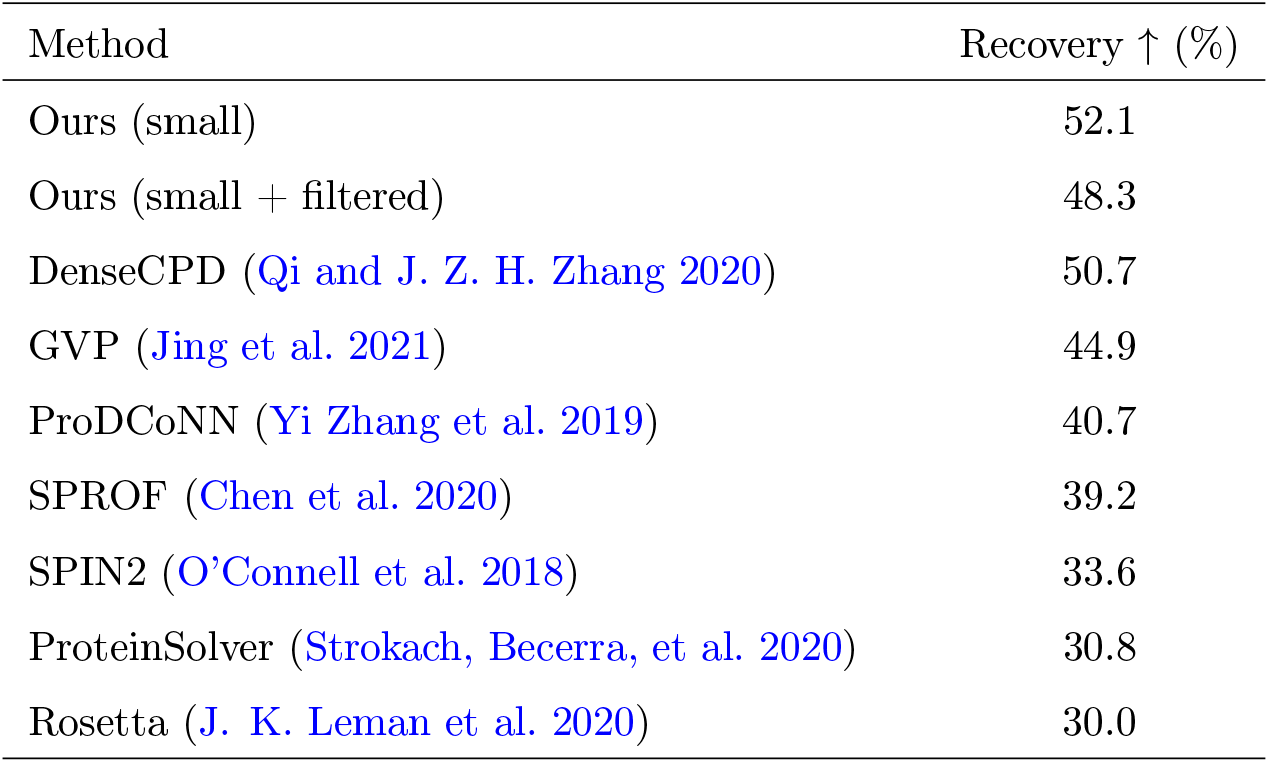
Results on TS50 Datasets. After filtering by sequence similarity, our small model achieves second best performance in NSR, behind 3DCNN-based method DenseCPD.

#### Results for CATH4.2 Dataset

For evaluating our model on CATH4.2 test chains, we trained a separate model using CATH4.2 chains for training (denoted *+ CATH4*.*2* in Figure S6). We also report results for our model trained on BC40 and evaluated on CATH4.2 test data. We note that there is potentially overlap between our training set and the CATH4.2 test chains.

**Figure S6:**
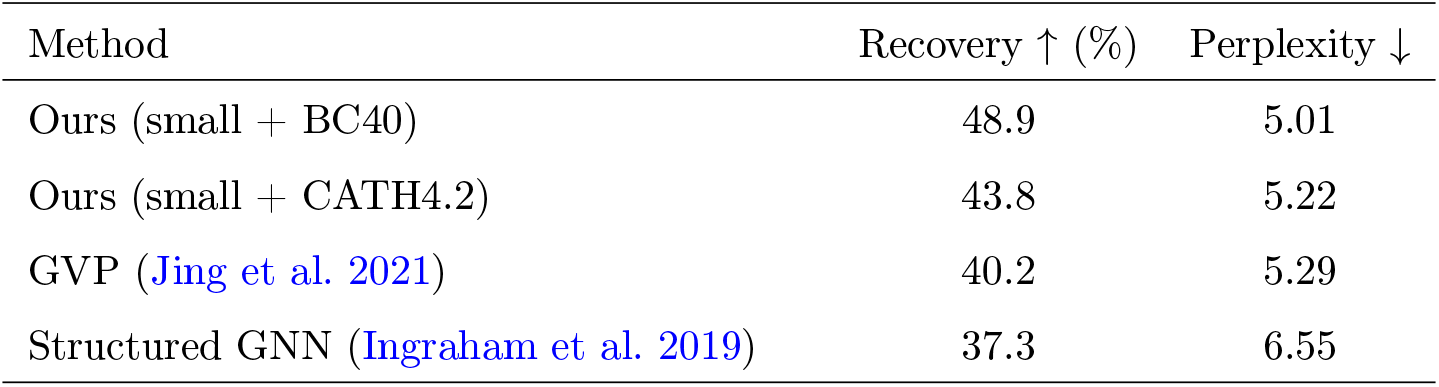
Results on CATH4.2 Dataset. Our small model trained on CATH4.2 data outperforms the structure transformer and GVP-GNN on the corresponding test set. Our model trained on BC40 achieves significantly higher recovery, but this may be caused by overlap between the train and test sets.

### S4 PDB Lists for Test Datasets

**Table S3:** List of targets in each test dataset.

| Dataset | Targets |
| --- | --- |
| CASP13 | T0949, T0950, T0951, T0953s1, T0953s2, T0954, T0955, T0957s1, T0957s2, T0958, T0959, T0960, T0961, T0962, T0963, T0964, T0965, T0966, T0967, T0968s1, T0968s2, T0969, T0970, T0971, T0973, T0974s1, T0974s2, T0975, T0976, T0977, T0978, T0979, T0980s1, T0980s2, T0981, T0982, T0983, T0984, T0985, T0986s1, T0986s2, T0987, T0988, T0989, T0990, T0991, T0992, T0993s1, T0993s2, T0994, T0995, T0996, T0997, T0998, T1000, T1001, T1002, T1003, T1004, T1005, T1006, T1008, T1009, T1010, T1011, T1013, T1014, T1015s1, T1015s2, T1016, T1016_A, T1017s1, T1017s2, T1018, T1019s1, T1019s2, T1020, T1021s1, T1021s2, T1021s3, T1022s1, T1022s2 |
| CASP14 | T1024, T1025, T1026, T1027, T1028, T1030, T1031, T1032, T1033, T1034, T1035, T1036s1, T1037, T1038, T1039, T1040, T1042, T1043, T1045s1, T1045s2, T1046s1, T1046s2, T1047s1, T1047s2, T1048, T1049, T1052, T1053, T1054, T1055, T1056, T1057, T1058, T1060s2, T1060s3, T1062, T1064, T1065s1, T1065s2, T1067, T1068, T1070, T1072s1, T1073, T1074, T1078, T1079, T1080, T1082, T1083, T1084, T1087, T1088, T1089, T1090, T1091, T1092, T1093, T1094, T1095, T1096, T1098, T1099, T1100 |
| TS50 | 1eteA 1v7mV 1yllA 3pivA 1or4A 2i39A 4gcnA 1bvyF 3on9A 3vjzA 3nbkA 3l4rA 3gwiA 4dkcA 3so6A 3lqcA 3gknA 3nngA 2j49A 3fhkA 2va0A 3hklA 2xr6A 3ii2A 2cayA 3t5gB 3ieyB 3aqgA 3q4oA 2qdlA 3ejfA 3gfsA 1ahsA 2fvvA 2a2lA 3nzmA 3e8mA 3k7pA 3ny7A 2gu3A 1pdoA 1h4aX 1dx5I 1i8nA 2cviA 3a4rA 1lpbA 1mr1C 2xcjA 2xdgA. |
| CATH4.2 | See (Ingraham et al. 2019), available here |

### Footnotes

1 Results obtained for DenseCPD have residue probabilities truncated to 3 significant digits. To compute perplexity, we replace probabilities of 0 with 10^−3^. Consequently, the perplexities obtained for DenseCPD in Table 1 serve as a lower bound on the true perplexity.

## Notes

### Competing Interest Statement

The authors have declared no competing interest.

